# PIKfyve deficiency in myeloid cells impairs lysosomal homeostasis in macrophages and promotes systemic inflammation in mice

**DOI:** 10.1101/610683

**Authors:** Sang Hee Min, Aae Suzuki, Lehn Weaver, Jessica Guzman, Liang Zhao, Francina Gonzalez, Lynn A. Spruce, Steven H. Seeholzer, Edward Behrens, Charles S. Abrams

**Author notes:** Department of Medicine, University of Michigan, Ann Arbor, MI 48109. **Corresponding Author:** Sang Hee Min.

## Abstract

Macrophages are professional phagocytes that are essential for host defense and tissue homeostasis. Proper membrane trafficking and degradative functions of the endolysosomal system is known to be critical for the function of these cells. We have found that PIKfyve, the kinase that synthesizes the endosomal phosphoinositide PI(3,5)P2, is an essential regulator of lysosomal biogenesis and degradative functions in macrophages. Genetically engineered mice lacking PIKfyve in their myeloid cells (*PIKfyve^fl/fl^ LysM-Cre*) develop diffuse tissue infiltration of foamy macrophages, hepatosplenomegaly, and systemic inflammation. PIKfyve loss in macrophages causes enlarged endolysosomal compartments and impairs the lysosomal degradative function. Moreover, PIKfyve deficiency increases the cellular levels of lysosomal proteins. Although PIKfyve deficiency reduced the activation of mTORC1 pathway and was associated with increased cleavage of TFEB proteins, this does not translate into transcriptional activation of lysosomal genes, suggesting that PIKfyve modulates the abundance of lysosomal proteins by affecting the degradation of these proteins. Taken together, our study shows that PIKfyve modulation of lysosomal degradative activity and protein expression is essential to maintain lysosomal homeostasis in macrophages.

## Introduction

Lysosomes are acidic organelles that are essential for the degradation of macromolecules delivered by endocytosis, phagocytosis and autophagy (1). Lysosomal degradation requires the hydrolytic enzymes and lysosomal membrane proteins that are continuously synthesized in the endoplasmic reticulum (ER), trafficked to the trans-Golgi network (TGN) and then sorted to the endo-lysosomal system (2). During this synthetic process, some lysosomal enzymes undergo a series of modifications that include cleavage of signal peptides in the ER, glycosylation by the addition of mannose-6-phosphate in the Golgi apparatus, and proteolytic processing of inactive zymogens into enzymatically active mature enzymes. Thus, proper functioning and homeostasis of lysosomes critically depend on the intracellular trafficking.

Intracellular trafficking events are modulated by several signaling molecules, which include members of the phosphoinositide family (3–5). In particular, the phosphatidylinositol (3,5)-bisphosphate [PI(3,5)P2] is critical for the trafficking along the endolysosomal system (6–8) and has been implicated in lysosome biogenesis and autophagy (9–12). PI(3,5)P2 is synthesized on the membranes of late endosomes and lysosomes (13, 14) by the lipid kinase, PIKfyve (phosphoinositide kinase, FYVE finger containing) (15–17). The physiological functions of PIKfyve have been recently elucidated in genetically engineered mice. PIKfyve-null mice are embryonic lethal, which indicates a critical role of PIKfyve during development (18, 19). A hypomorphic PIKfyve mouse model is viable, but dies early due to defects in multiple organs including neural tissues, heart, lung, kidney, thymus and hematopoietic system (20). Subsequently, conditional PIKfyve knockout mice were developed using the cre-lox system, and demonstrated the essential roles of PIKfyve in specific tissues (19, 21–26). In particular, we previously showed that mice with platelet-specific deletion of PIKfyve have impaired lysosomal homeostasis and develop aberrant inflammatory and prothrombotic responses (22). Similarly, intestine-specific PIKfyve knockout mice develop defective polarization of epithelial cells, which leads to a severe inflammatory bowel disease (19). However, recently generated myeloid-specific PIKfyve knockout mice did not show any abnormalities in the resident macrophages in spleen, liver or bone marrow, but affected only some populations of alveolar macrophages in the lung (25). We were surprised by this latter observation given the known importance of lysosome function within professional phagocytic cells, and published effect of PIKfyve inhibitors on innate immunity (27, 28). Therefore, we chose to revisit the role of PIKfyve in myeloid cells using a myeloid-specific PIKfyve knockout mouse that was generated using our previously reported PIKfyve flox mice (22).

We found that PIKfyve deficiency in myeloid cells leads to proliferation of granulocytes and monocytes associated with elevation of inflammatory cytokines, and that PIKfyve is a critical regulator of the morphology, degradative activity and protein turnover of the endolysosomal system in macrophages. In summary, our study demonstrates that PIKfyve is essential for maintaining lysosomal homeostasis and function in macrophages.

## Results

### Generation and validation of mice lacking PIKfyve in myeloid lineage

Previously reported myeloid-specific PIKfyve knockout mice were generated using mice with lox P sites flanking exon 5 of the PIKfyve gene (25). Since this targeting approach can sometime produce truncated proteins from cryptic start sites and lead to hypomorphic phenotypes, we used our previously published mice with lox P sites flanking exons 37 and 38 (corresponding to the kinase domain) of the PIKfyve gene (PIKfyve^fl)^ (22) to breed with mice expressing the recombinase Cre under the LysM promoter and generate *PIKfyve^fl/fl^ LysM-Cre* mice (Fig. 1A). To validate the tissue specificity of LysM-Cre, *PIKfyve^fl/fl^ LysM-Cre* mice were bred with Cre-dependent YFP reporter mice to obtain *PIKfyve^fl/fl^ LysM-Cre Rosa26YFP* mice. The peripheral blood of *PIKfyve^fl/fl^ LysM-Cre Rosa26YFP* mice were analyzed for YFP expression by flow cytometry, which demonstrated that the LysM-Cre promotor induced Cre expression in about 80% of the monocytes, in about 90% of the neutrophils, and only in about 1% - 5% of the circulating B cells and T cells (Fig. 1B). Primary tissue macrophages were isolated from the spleen or bone marrow using F4/80 antibody by immunomagnetic separation, and were analyzed for the expression of PIKfyve. PIKfyve mRNA expression was significantly reduced in the macrophages of *PIKfyve^fl/fl^ LysM-Cre* mice compared to *wild-type (WT) Lysm-Cre* mice as determined by qRT-PCR analysis (Fig. 1C). Furthermore, PIKfyve protein expression in macrophages was partially reduced in the *PIKfyve^fl/+^ LysM-Cre* mice as compared to *WT LysM-Cre* mice and completely undetectable in the *PIKfyve^fl/fl^ LysM-Cre* mice (Fig. 1D). Given that our gene targeting removed exons which encoded essential components of the PIKfyve kinase domain, but still allowed expression of a truncated mRNA, this finding suggests that the resulting truncated PIKfyve mRNA is likely unstable and undergoes degradation, leading to the complete loss of PIKfyve protein in their macrophages.

**Figure 1:**
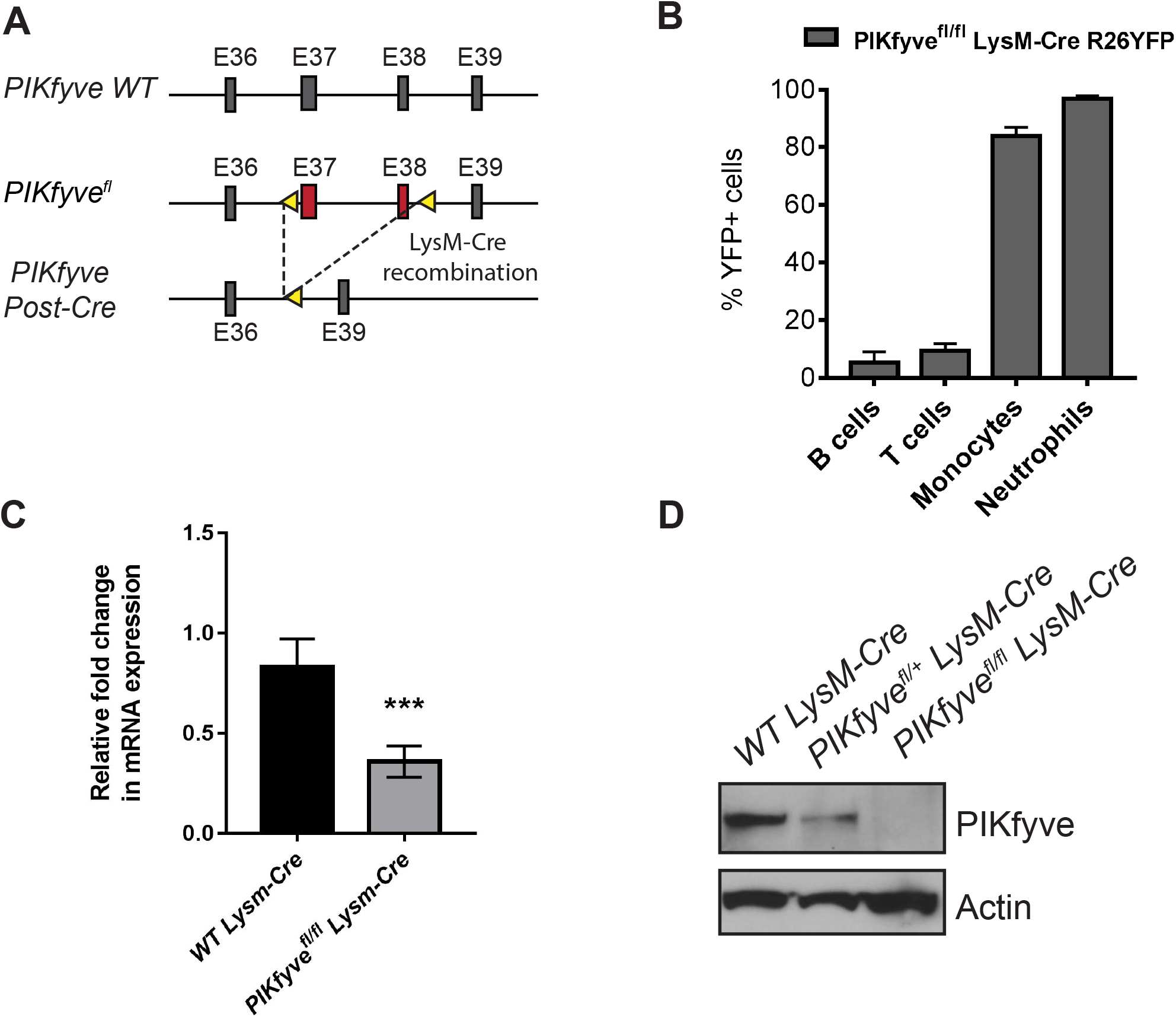
Generation and validation of mice lacking PIKfyve in myeloid cells. (A) Schematic depicting the genetic targeting of *PIKfyve. PIKfyve-floxed* alleles (*PIKfyve^fl^*) were generated by targeting the exons 37 and 38 with lox P sites (yellow arrows). *PIKfyve^fl^* mice were crossed with *LysM-Cre* mice to generate a myeloid-specific homologous recombination of *PIKfyve^fl^*. (B) Flow cytometry analysis of percentage of YFP expressing cells in the peripheral blood of *PIKfyve^fl/fl^ LysM-CreR26 YFP* mice (*n* = 4 mice). (C) qRT-PCR analysis of PIKfyve gene expression relative to GAPDH in the F4/80+ spleen macrophages (*n* = 3 mice). (D) Immunoblotting analysis of PIKfyve protein expression in the F4/80+ spleen macrophages. \**P*<0.05, \*\*\**P*<0.001, NS, *P*>0.05. All error bars indicate mean +/− s.e.m. Unpaired two-tailed Student’s t-test.

### PIKfyve ablation in myeloid cells causes tissue accumulation of vacuolated macrophages and promotes systemic inflammation

Previously reported myeloid-specific PIKfyve knockout mice did not develop any gross abnormalities (25). Although our *PIKfyve^fl/fl^ LysM-Cre* mice were born at the expected Mendelian frequency and displayed no discernible morphological abnormalities at birth, they developed progressive abdominal distention as they matured (Fig. 2A). Necropsy at different ages showed that *PIKfyve^fl/fl^ LysM-Cre* mice developed enlargement of their livers and spleens compared to their *WT LysM-Cre* littermates (Fig. 2B and Fig. 2C). Histological analysis of these organs revealed tissue accumulation of highly vacuolated macrophages (Fig. 2D). Immunophenotyping analysis of circulating leukocytes from *PIKfyve^fl/fl^ LysM-Cre* mice showed significantly increased numbers of neutrophils and monocytes, suggesting a systemic inflammatory response (Fig. 2E). Further analysis of plasma samples for cytokine profiling by multiplex assay revealed elevated levels of several pro-inflammatory and chemotactic cytokines including IL-6, IL-20, CCL-4, CCL-19, CXCL-9, CXCL-10, eotaxin and TIMP-1 (Fig. 2F) in *PIKfyve^fl/fl^ LysM-Cre* mice compared to *WT LysM-Cre* littermates. Together, these findings demonstrate that PIKfyve deficiency can result in a pathologic process that is reminiscent of lysosomal storage disorders and is associated with systemic inflammation.

**Figure 2:**
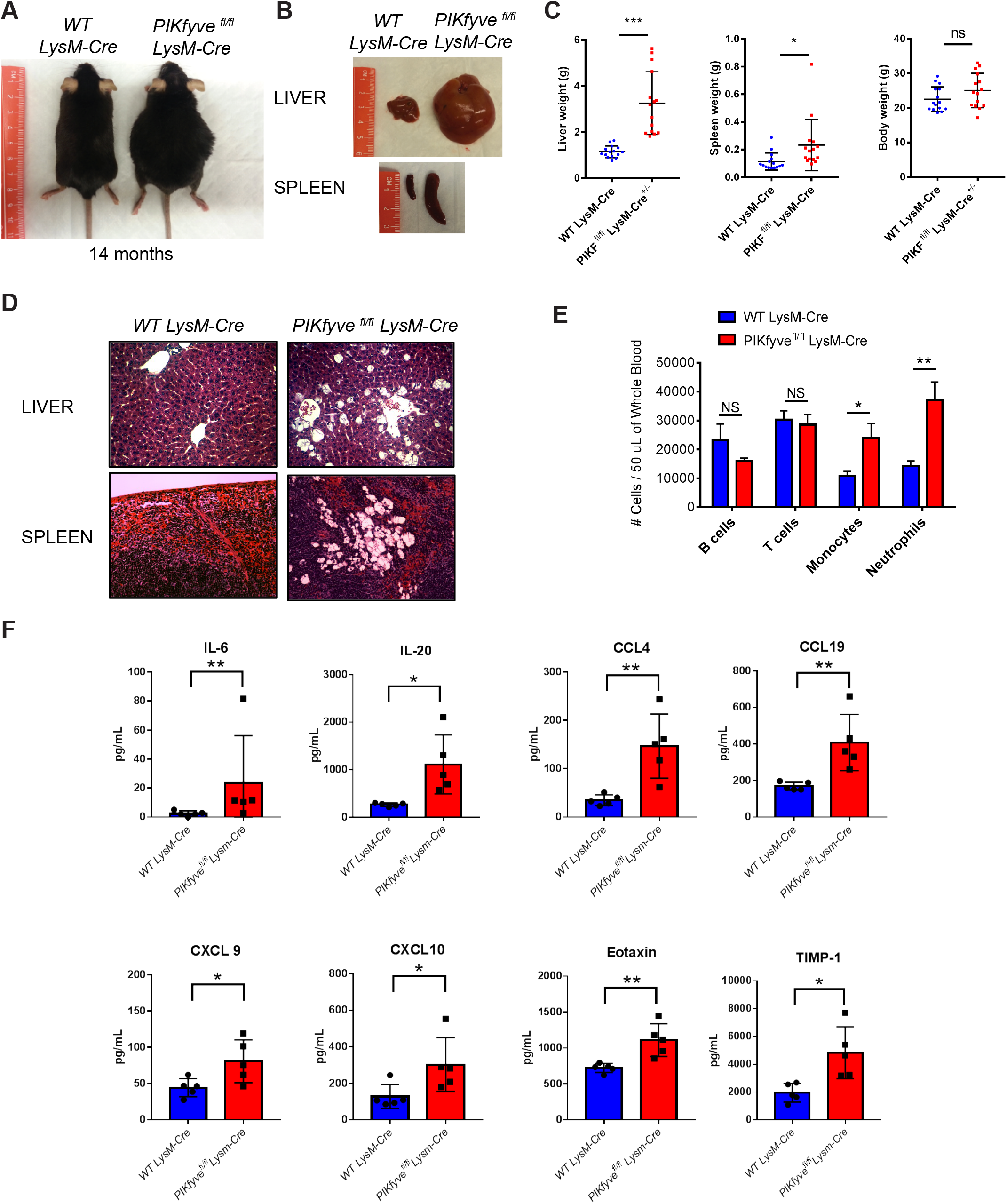
*PIKfyve^fl/fl^ LysM-Cre* mice develop features of a lysosomal storage disorder. (A) General appearance of mice at about 14 months of age. Note the characteristic abdominal distention in the *PIKfyve^fl/fl^ LysM-Cre* mouse. (B) Representative images of the liver and spleen of mice at 14 months of age, illustrating the marked hepatosplenomegaly in the *PIKfyve^fl/fl^ LysM-Cre* mouse. (C) Average weights of the body, liver, and spleen from mice at 8-35 weeks of age (*n* = 15 per group). (D) Representative images of tissue sections of the liver and spleen stained with hematoxylin and eosin. Note the tissue accumulation of engorged cells with translucent cytoplasmic vacuoles in the *PIKfyve^fl/fl^ LysM-Cre* mouse. (E) Flow cytometry analysis of the numbers of B lymphocytes, T lymphocytes, monocytes and neutrophils in the peripheral blood of mice at age of 4-20 weeks of age (*n* = 7 for *WT LysM-Cre* and *n* = 11 for PIKfyve^*fl/fl*^ *LysM-Cre*). (F) Cytokine analysis by multiplex array of plasma samples of mice at 8-16 weeks of age (*n* = 5 per group). \**P*<0.05, \*\*\**P*<0.001, NS, *P*>0.05. All error bars indicate mean +/− s.e.m. Unpaired two-tailed Student’s t-test.

### PIKfyve regulates lysosomal structure and proteolytic function in macrophages

To investigate the effect of PIKfyve deficiency in macrophages, we first examined the morphology of macrophages isolated from the bone marrow or spleens of *PIKfyve^fl/fl^ LysM-Cre Rosa26YFP* mice. We confirmed that macrophages isolated using F4/80 antibody were YFP+ indicating LysM-Cre expression in F4/80+ macrophages (Fig 3A). As expected, PIKfyve-null macrophages displayed cytoplasmic vacuolation (Fig 3A), which was similar to the vacuolation that has been previously reported in other PIKfyve-null cells (19, 20, 22). The enlarged cytoplasmic vacuoles in PIKfyve-null macrophages expressed LAMP1, which is a marker of late endosomes or lysosomes (Fig 3B). Together, these data confirm that PIKfyve is necessary to maintain the endolysosomal morphology.

**Figure 3:**
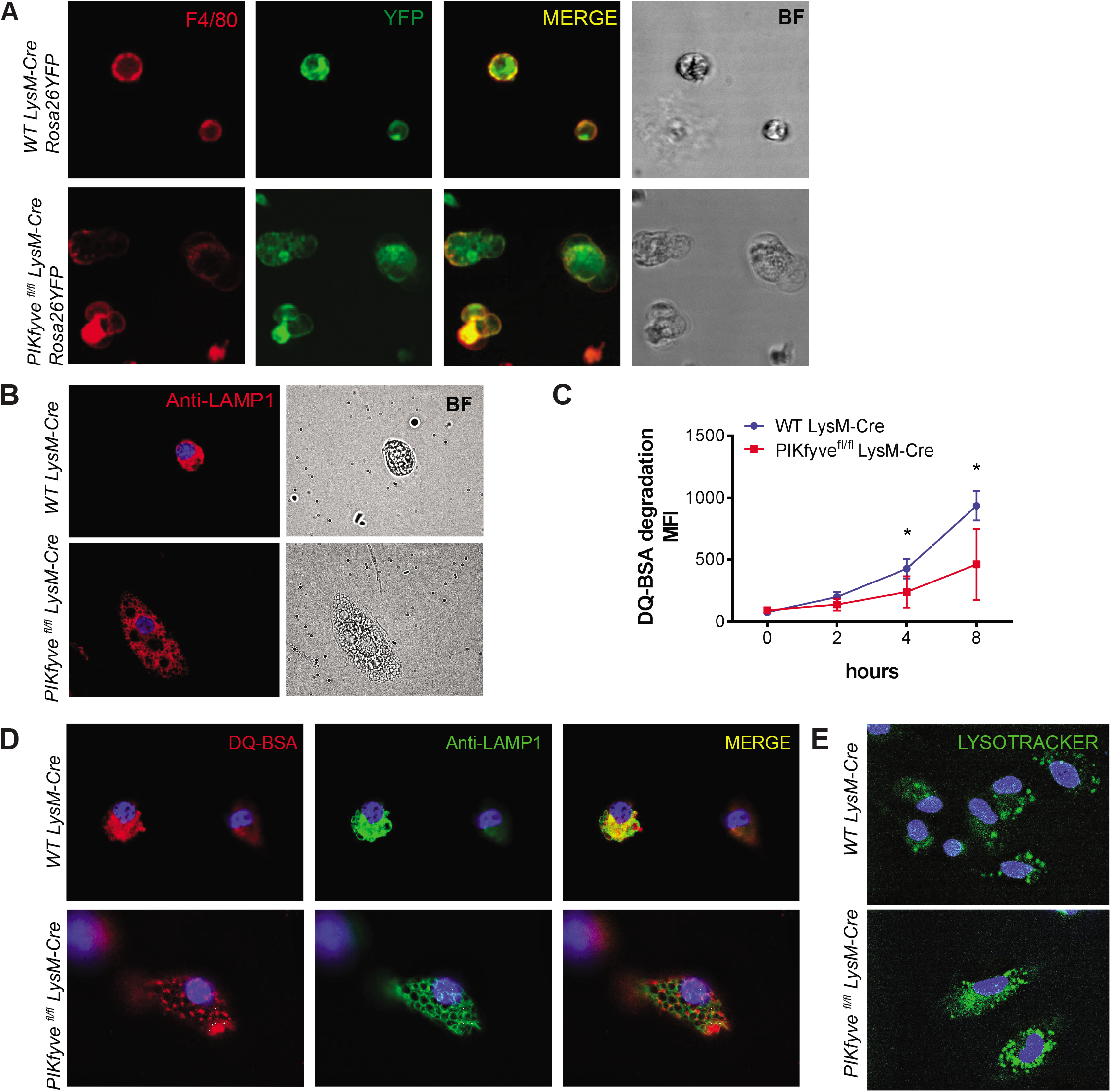
PIKfyve regulates lysosomal morphology and degradative function in macrophages. (A) Live cell fluorescence microscopy images of spleen macrophages isolated and stained with F4/80 antibody. The intrinsic YFP expression is driven by *LysM-Cre*. Note the presence of multiple cytoplasmic vacuoles of various sizes in the macrophages of *PIKfyve^fl/fl^ LysM-Cre Rosa26YFP* mouse. (B) Immunofluorescence images of bone marrow derived macrophages stained with anti-LAMP1 antibody. (C) Proteolytic DQ-BSA degradation over eight hours in the F4/80+ spleen macrophages as measured by increasing fluorescence of quenched dye on a spectrophotometer. (D) Immunofluorescence images of bone marrow derived macrophages incubated with DQ-BSA for one hour and co-stained with anti-LAMP-1 antibody. (E) Images of bone marrow derived macrophages incubated with LysoTracker for 20 minutes. Note the accumulation of LysoTracker in the acidic endolysosomes. \**P*<0.05, NS, *P*>0.05. All error bars indicate mean +/− s.e.m. Analysis was done using an unpaired two-tailed Student’s t-test.

We next investigated whether PIKfyve was essential for the degradative function of macrophage lysosomes. Lysosomal proteolytic degradation was determined using self-quenched DQ-BSA, which is a protease substrate that is taken up by endocytosis and emits fluorescence upon proteolytic degradation within acidic compartments such as late endosomes and lysosomes. Proteolysis of DQ-BSA was detected by two independent techniques. First, we analyzed the proteolytic degradation of DQ-BSA in the lysates of macrophages via spectrophotometry. PIKfyve-null macrophages had significantly impaired ability to proteolytically degrade DQ-BSA (Fig. 3C). For the second method to determine the effect of PIKfyve on lysosomal proteolytic degradation in live cells, we analyzed the cellular localization of DQ-BSA degradation. As anticipated, WT macrophages displayed a robust ability to catabolize DQ-BSA within LAMP1 demarcated compartments (Fig. 3D). In contrast, PIKfyve-null macrophages showed undetectable proteolytic degradation of DQ-BSA within their enlarged LAMP1-positive late endosomes and lysosomes. Together, these results demonstrate the critical role of PIKfyve in the ability of macrophages to degrade proteins within their lysosomes.

As a low pH is necessary for normal proteolytic function of lysosomal enzymes, we further analyzed whether the absence of lysosomal proteolysis within PIKfyve-null macrophages was due to a necessary role for PIKfyve in lysosomal acidification. This process was analyzed using Lyso Tracker, a fluorescent dye that accumulates in acidic compartments such as lysosomes. We found that Lyso Tracker accumulated in the enlarged cytoplasmic vacuoles in the macrophages of *PIKfyve^fl/fl^ LysM-Cre* mice (Fig. 3E). This demonstrates that PIKfyve is not required for the acidification of endolysosomal compartments.

### PIKfyve modulates lysosomal protein abundance independently of transcription

Given the importance of PIKfyve in lysosomal morphology and function, we next investigated the role of PIKfyve in lysosomal biogenesis. First, we examined the abundance of lysosomal and autophagy-related proteins in the F4/80+ macrophages from *WT LysM-Cre* mice and *PIKfyve^fl/fl^ LysM-Cre* mice by immunoblotting analysis. Compared to WT macrophages, PIKfyve-null macrophages showed increased levels of lysosomal and autophagy-related proteins such as LAMP1, procathepsin D, cathepsin D, and LC3 (Fig. 4A).

**Figure 4:**
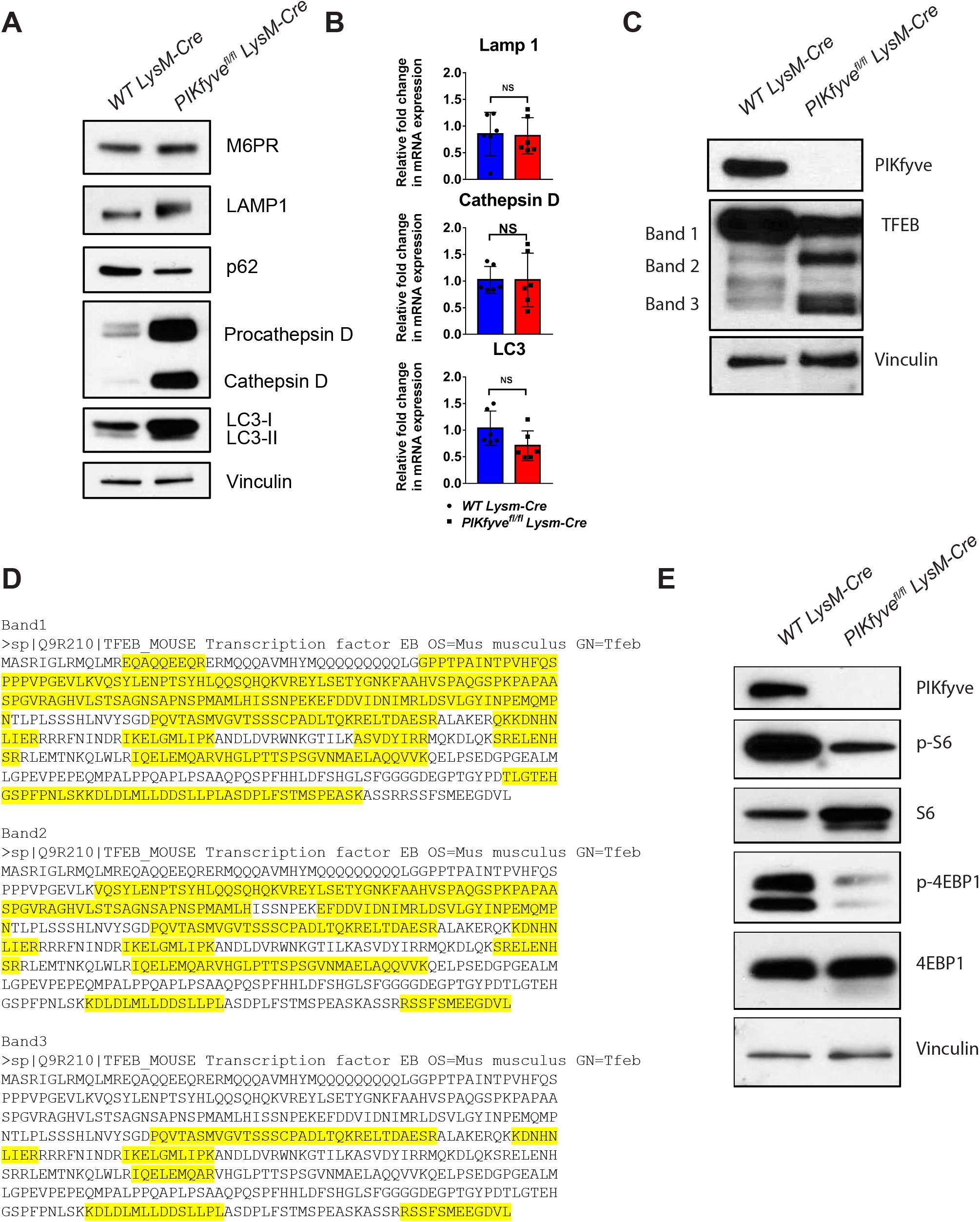
Effects of PIKfyve deficiency in lysosomal biogenesis and protein expression. (A) Immunoblot analysis of the F4/80+ spleen macrophages for lysosomal and autophagy-related proteins: M6PR, LAMP1, p62, procathepsin D, cathepsin D, LC3-I, and LC3-II. Vinculin was used as a loading control. (B) qRT-PCR analysis of cathepsin D, LAMP1 and LC3 relative to GAPDH in the F4/80+ spleen macrophages (*n* = 3 mice). (C) Immunoblot of the F4/80+ spleen macrophages probed with the TFEB antibody. TFEB protein bands 1, 2, and 3 were cut for mass spectrometry analysis. (D) Proteomic analysis of TFEB protein bands 1, 2 and 3 in the figure 4C. Highlighted in yellow are the tryptic peptides identified in each band by mass spectrometry analysis. Uniprot Q3UKG7 was used for the amino acid sequence of murine TFEB. (E) Immunoblot analysis of the F4/80+ spleen macrophages for PIKfyve, p-S6, S6, p-4EBP1, and 4EBP1. Probing for vinculin was used as the loading control. All error bars indicate mean +/− s.e.m. Analysis was done using an unpaired two-tailed Student’s t-test.

Lysosomal function and biogenesis is closely regulated by the transcription factor TFEB, which is often referred to as the “master regulator” of the lysosomal gene network (29). Based on this premise, we hypothesized that the elevated lysosomal proteins in PIKfyve-null macrophages is secondary to activation of TFEB promoting transcriptional upregulation of lysosomal genes. In contrast to our prediction, the quantity of expressed mRNA for LAMP1, cathepsin D or LC3 was not significantly higher in the PIKfyve-null macrophages compared to the WT macrophages by qRT-PCR analysis (Fig. 4B). Intriguingly, we found that PIKfyve-null macrophages had reduced levels of full-length TFEB of about ~60KD (band 1, Fig. 4C) but increased amounts of the shorter variants of the TFEB protein that were between 40-50KD (bands 2 and 3, Fig. 4C). Further analysis of the different forms of TFEB by mass spectrometry confirmed that these variants were N-terminal truncated variants of TFEB (Fig. 4D), although the presence and significance of TFEB truncation variants have not been previously reported. We propose that these truncated variants are likely inactive forms of TFEB since their presence were not associated with upregulated lysosomal gene expression.

Previous studies showed that the activity of TFEB is regulated primarily by mTORC1-mediated phosphorylation (30–32). Thus, we examined the effect of PIKfyve ablation in mTORC1 activation. This was done by analyzing the effect of the PIKfyve-null mutation on the mTORC1 substrates phospho-S6 and phospho-4EBP1. Interestingly, PIKfyve-null macrophages had decreased levels of phospho-S6 and phospho-4EBP1 compared to WT macrophages by immunoblotting (Fig. 4E), which indicated reduced activity of mTORC1. Together, these findings suggest that the activity of mTORC1 and TFEB are disconnected in the PIKfyve-null macrophages. Furthermore, our findings suggest that PIKfyve deficiency leads to the accumulation of lysosomal proteins likely from reduced degradation of lysosomal proteins and not from TFEB-mediated transcriptional activation of lysosomal genes.

## Discussion

In this study, using mice genetically engineered to lack PIKfyve in their myeloid cells, we found that PIKfyve is an essential regulator of the lysosomal morphology, degradative activity and protein abundance in macrophages. Although our findings are consistent with the phenotype seen in cells exposed to the PIKfyve inhibitor (apilimod) (27, 28) and genetic loss of function mutations in embryonic stems cells (18, 20), these findings are different from the phenotype of the previously reported myeloid-specific PIKfyve knockout mice (25), in which PIKfyve was dispensable for most tissue-resident macrophages except for alveolar macrophages in the lung. There are three possible explanations for why we find PIKfyve so essential for macrophage biology while another targeting strategy does not: 1) Our mice were generated in a pure C57BL/6 background whereas Kawasaki et al. generated their mice by injection of ES cells from JM8/A3 strain into C57BL/6 background mice; 2) While Kawasaki et al. targeted the exon 5 of PIKfyve gene, we targeted the exons 37 and 38 of the PIKfyve gene. It is possible that our targeting approach which removes critical components of the kinase domain may result in a truncated form that functions as a dominant negative gene. However, the absence of a phenotype in the heterozygous *PIKfyve^fl/+^ LysM-Cre* mice argues against this possibility; and 3) It is possible that the targeting strategy utilized by Kawasaki et al., which removes the start codon still permits expression from a cryptic start site that generates a small amount of truncated protein that is catalytically active and produces a hypomorph. Such a residual minor amount of PIKfyve protein might be difficult to detect but still could be sufficient to generate a small amount of PI(3,5)P2 that attenuates the macrophage phenotype.

With the exception of the above referenced publication, studies using genetic or pharmacologic ablation of PIKfyve in mammals or of its orthologue fab1p in yeast, consistently demonstrate the development of enlarged endolysosomal compartments (33–36). Interestingly, fab1-null yeast have impaired acidification of their enlarged vacuoles (33), whereas our PIKfyve-impaired macrophages, have preserved lysosomal acidification. Consistent with prior studies using PIKfyve inhibitors (27), we found that mammalian PIKfyve is dispensable for the regulation of lysosomal pH, since the enlarged late endosomes and lysosomes maintained the acidification in the PIKfyve-null macrophages. Additionally, our data showing the absence of DQ-BSA cleavage within the LAMP1-demarcated compartments, now demonstrates that despite normal acidification, the enlarged late endosomes and lysosomes in PIKfyve-null macrophages have impaired proteolytic activity. Curiously, proteolysis still occurred in some compartments that were not demarcated by LAMP1. This suggests that DQ-BSA is internalized and proteolyzed, perhaps in earlier endosomal compartments, but it cannot be degraded in the downstream compartments, such as in the late endosomes and lysosomes.

The defective degradation within the lysosomes could be explained by the following. First, proteins that are targeted for degradation could require PIKfyve in order for them to be transported to the late endosomes or lysosomes. Second, PIKfyve could be necessary for the trafficking of critical degradative proteases to late endosomes or lysosomes. Lastly, PIKfyve could be required for an essential step in the post-translational processing of lysosomal degradative proteases that is required for their enzymatic activity. Consistent with several observations in models of PI(3,5)P2 deficiency (20, 22, 37), PIKfyve-null macrophages have disproportionally increased levels of Procathepsin D, which is unprocessed form of Cathepsin D. These findings suggest that PIKfyve is critical for the processing and maturation of lysosomal proteins.

Our study demonstrates that PIKfyve is necessary for the regulation of expression levels of lysosomal proteins. We initially hypothesized that this would be driven by activation of TFEB and consequently increased transcriptional expression of lysosomal genes. In contrast to this hypothesis, we observed that PIKfyve deficiency is instead associated with a decreased activation of mTORC1 as well as decreased expression and activation of TFEB. In spite of increased amounts of lysosomal proteins within PIKfyve-null macrophages, the expression of lysosomal genes was not elevated in PIKfyve-null macrophages. Together, these findings suggest that PIKfyve does not directly modulate lysosomal gene expression, but instead is important for the turnover of lysosomal proteins. However, the exact mechanism of PIKfyve modulating lysosomal protein turnover still remains to be elucidated.

In conclusion, our study shows that PIKfyve is essential to maintain lysosomal homeostasis in macrophages by regulating the morphology, trafficking, degradative function and protein expression in the lysosome.

## Materials and Methods

### Mice

Mice expressing PIKfyve floxed alleles (*PIKfyve^fl^*) with the exons 37 and 38 of the PIKfyve gene flanked by loxP sites were generated as previously described (22). To generate myeloid-cell specific PIKfyve-deficient mice, *PIKfyve^fl^* mice were crossed to mice expressing Cre recombinase under the control of the endogenous lysozyme 2 promoter (LysM-Cre) (B6.129P2-*Lyz2^tm1(cre)Ifo^*/J - stock # 004781, Jackson laboratory). The resulting *PIKfyve^fl/fl^ LysM-Cre* mice were crossed to a Cre reporter strain that expresses EYFP upon cre-mediated recombination (B6. Cg-*Gt(ROSA)26Sor^tm3(CAG-EYFP)Hze^*/J - stock #007903, Jackson laboratory). For all studies, both female and male mice were used. All mice were maintained on standard chow and tap water in pathogen-free conditions. All animal procedures were approved by and performed in accordance with the Institutional Animal Care and Use Committee at the University of Pennsylvania.

### PCR Genotyping

Genomic DNA was isolated from mouse tail biopsies for PCR genotyping. Genotyping for PIKfyve^fl^ was performed as previously described(22). Briefly, LoxP integration in the Intron 36 was identified with 5′ - CCATTGCCTGGCTTAGAACAGAG -3′ and 5′ - GAACTCTCCCGCGTAGTACAGC -3′ primers. LysM-Cre was identified with mutant primer 5’ - CCC AGA AAT GCC AGA TTA CG -3’, common primer 5’ -CTT GGG CTG CCA GAA TTT CTC -3’, and WT primer 5’ - TTA CAG TCG GCC AGG CTG AC -3’. Rosa26/EYFP transgene was identified with WT F 5’ -AAG GGA GCT GCA GTG GAG TA-3’ WT R 5’- CCG AAA ATC TGT GGG AAG TC-3’ mutant F 5’- ACA TGG TCC TGC TGG AGT TC-3’ and mutant R 5’- GGC ATT AAA GCA GCG TAT CC – 3’.

### Whole Blood WBC Analysis

Whole blood (50 μL) was obtained by retro-orbital bleed and red blood cells were lysed in ACK lysing buffer (Lonza). The remaining cells were stained with Live Dead Aqua (Life Technologies) and incubated with cell culture supernatants from anti-CD16/32 (clone 24G2) expressing cells to block non-specific binding to the Fc receptor. Cells were subsequently stained with fluorophore-labeled monoclonal antibodies (mAb) to the following antigens: CD3 (clone 145-2C11), CD19 (clone 6D5), CD115 (clone AFS98), CCR2 (clone 475301), Ly6C (clone HK1.4), and Ly6G (clone 1A8) that were obtained from BD Pharmingen, Biolegend, or from R&D Systems. Samples were analyzed on a MacsQuant flow cytometer (Miltenyi) and flow cytometric analysis was performed using FlowJo software (Tree Star).

### Immunomagnetic Isolation of macrophages from the spleen or bone marrow

The protocol was adapted from Stemcell Technologies. Bone marrow was flushed from mice femurs, aspirated with a syringe, and filtered with a cell strainer to derive a single cell suspension. Spleens were dissected from the abdominal cavity of mice and filtered through a 50um nylon strainer to make single cell suspension of splenocytes. Red blood cells were lysed with ACK buffer. Next, the cells were blocked with FcR block (Miltenyi Biotec 130-092-575), incubated with anti-F4/80 antibodies conjugated with FITC/PE (Miltenyi Biotec 130-102-988, 130-102-943), and incubated in EasySep FITC/PE Positive Selection kit which labels FITC/PE epitopes with magnetic beads (Stemcell Technologies 18557, 18555). Each step was followed by a wash and spin. The prepared cells (in 15 mL centrifuge tubes) were placed in EasySep magnets (Stemcell Technologies 18001) which retained labeled cells during washes. After several washes to remove unlabeled cells, the tubes were removed from the magnets and the isolated cells were collected.

### Macrophage Culture

To generate bone marrow derived macrophages (BMDM), bone marrow cells were extracted from femurs and tibias of mice at 8-12 weeks of age, and cultured in DMEM/F12 (Thermo Fisher). The cells were supplemented with 10% FBS, 1% penicillin and streptomycin, and recombinant mouse M-CSF (Calbiochem) at a final concentration of 10ng/mL for seven days. Supernatant cells were discarded, and BMDM were harvested from dishes by adding Accutase (Sigma Aldrich A6964) and washing with DMEM.

### Histochemistry

Tissues were harvested, fixed overnight in 10% formalin, paraffin embedded, and sectioned. Paraffin sections were deparaffinized, and stained with hematoxylin and eosin.

### Real-time quantitative PCR

RNA was isolated from F4/80+ macrophages using Illustra RNA Spin Mini kit (GE Healthcare). cDNA was made with High Capacity cDNA kit (Thermo Fisher Scientific). 100-400ng of cDNA was used for qPCR. Primers and probes were acquired from Thermo Fisher Scientific: Gapdh (Mm99999915_g1); PIKfyve (Mm00440793_m1); LC3 (Mm00458724_m1); Cathepsin D (Mm00515586_m1) and Lamp1 (Mm00495262_m1). Samples were run on a Step One Plus q-PCR instrument (Applied Biosystems) and analyzed using delta-delta Ct method to calculate fold change. All samples were first normalized to Gapdh and then compared to WT controls.

### Immunoblotting

Tissues or cells were harvested and homogenized in RIPA buffer that was supplemented with a protease inhibitor cocktail (Sigma-Aldrich) and a phosphatase inhibitor cocktail (Thermo Fisher Scientific). The protein concentrations were measured by the BCA Protein Assay (Thermo Fisher Scientific). The protein samples were analyzed by novex NuPage SDS-PAGE gradient gels under reducing conditions (Invitrogen), and then transferred onto the polyvinylidene difluoride membrane (Invitrogen). The membrane was blotted with the indicated primary antibodies against: PIKfyve (Sigma-Aldrich; 1:400), M6PR (Abcam; 1:1000), LAMP1 (Developmental Studies Hybridoma Bank; 1:2000), p62 (Cell Signaling Technology; 1:1000), Cathepsin D (Calbiochem 1:1000), LC3 (cell signaling technology; 1:1000), TFEB (Bethyl; 1:2000), Phospho S6 (Cell Signaling Technology; 1:1000), S6 (Cell Signaling Technology1:1000), Phospho 4EBP1 (Cell Signaling Technology; 1:1000), 4EBP1 (Cell Signaling Technology; 1:1000), β-actin (Cell Signaling Technology; 1:2000) and Vinculin (Santa Cruz Biotechnology; 1:1000). The following horseradish peroxidase-conjugated secondary antibodies were used: anti-rabbit (GE Healthcare; 1:3000), anti-rabbit (Cell Signaling Technology; 1:1000), anti-rat (Santa Cruz Biotechnology; 1:5000), anti-goat (Santa Cruz Biotechnology; 1:3000), and anti-mouse (Santa Cruz Biotechnology; 1:3000). Membranes were visualized with enhanced chemiluminescence substrate (GE Healthcare Life Sciences).

### Mouse cytokine multiplex assay

Plasma samples were obtained from mice and analyzed by Eve technologies (Calgary, Canada) using the by the mouse cytokine array / chemokine array 44-plex according to the manufacturer’s instructions. The following biomarkers were analyzed: Eotaxin, Erythropoietin, 6Ckine, Fractalkine, G-CSF, GM-CSF, IFNB1, IFNγ, IL-1α, IL-1β, IL-2, IL-3, IL-4, IL-5, IL-6, IL-7, IL-9, IL-10, IL-11, IL-12 (p40), IL-12 (p70), IL-13, IL-15, IL-16, IL-17, IL-20, IP-10 (CXCL10), KC, LIF, LIX, MCP-1, MCP-5, M-CSF, MDC, MIG (CXCL9), MIP-1α (CCL3), MIP-1β (CCL4), MIP-2 (CXCL2), MIP-3α (CCL20), MIP-3B (CCL19), RANTES, TARC, TIMP-1, TNFα, and VEGF.

### Immunofluorescence Microscopy

BMDM were grown on coverslips and fixed with cold MeOH/acetone, permeabilized with PBT (PBS and Triton X 0.1%), and blocked with Starting Block buffer T20 (Thermo FisherScientific). Slides were probed with primary antibody LAMP1 (Development Study Hybridoma Bank) overnight at 4°C. Slides were imaged with Leica DM6000, and images were deconvolved with Leica LAS Autoquant software.

### Live cell imaging with Lyso Tracker Green and DQ-Red BSA staining

Macrophages were grown overnight on glass chamber slides (Ibidi #80827) coated with 0.1% gelatin. For all live stain solutions and washes, a 10% FBS DMEM (w/o phenol red; Invitrogen) staining buffer was used. Macrophages were gently washed with staining buffer and labeled with live stains LysoTracker Green or DQ-Red BSA (Thermo Fisher Scientific L7526, D12051). To visualize lysosomes, LysoTracker Green (100 μM) was applied to each well for 15 minutes. To measure proteolytic activity, DQ-Red BSA (10 μg/mL) was added to each well for different time periods. For each stain, the wells were washed once with 10% trypan blue in staining buffer followed by a wash with staining buffer and then fixed with 4% paraformaldehyde. Images were acquired with Metamorph software on a spinning disk confocal microscope (Nikon Eclipse Ti-U) at the University of Pennsylvania Molecular Pathology & Imaging Core Service.

### Proteolysis Assay

BMDM were seeded into a 96-well plate and incubated overnight at 37°C. Cells were then incubated with PBS or DQ-Red BSA (Thermo Fisher Scientific) at a final concentration of 10 μg/mL and incubated at 37°C for 0, 2, 4, and 8 hours. Fluorescence of DQ-Red BSA was measured on a Molecular Devices spectrophotometer microplate reader at excitation 584 nm and emission 612 nm.

### Immunoprecipitation of TFEB

Isolated macrophages from the mouse spleen were lysed in RIPA (or in 1% NP40 in PBS) lysis buffer, containing a protease inhibitor cocktail (Roche) and sodium orthovanadate (2 mM). The lysates were precleared with unconjugated protein A agarose beads (Invitrogen). The precleared lysates were subjected to immunoprecipitation using anti-TFEB antibody (Bethyl Laboratories A303-673A) prebound to protein A agarose beads. The immunoprecipitated sample was eluted by adding Novex NuPAGE LDS Sample Buffer (Invitrogen) and Novex NuPAGE Sample Reducing Agent (Invitrogen), and heating at 95°C for 10 minutes. The sample was spun down at 500g x five minutes, and the supernatant was run onto 4%-12% Novex NuPAGE gel (Invitrogen). The gel was Coomassie blue stained and the corresponding lanes to TFEB bands were excised for digestion and mass-spectrometry analysis.

### In-gel digestion

Each sample was excised from the gel and cut into 1 mm^3^ cubes (38). They were destained with 50% methanol/1.25% acetic acid, reduced with 5 mM dithiothreitol (Thermo Fisher Scientific), and alkylated with 40 mM iodoacetamide (Sigma-Aldrich). Gel pieces were then washed with 20 mM ammonium bicarbonate (Sigma-Aldrich) and dehydrated with acetonitrile (Thermo Fisher Scientific). Trypsin (Promega; 5 ng/mL in 20 mM ammonium bicarbonate) was added to the gel pieces and proteolysis was allowed to proceed overnight at 37 °C. Peptides were extracted with 0.3% triflouroacetic acid (J.T.Baker), followed by 50% acetonitrile. Extracts were combined and the volume was reduced by vacuum centrifugation.

### Mass spectrometry analysis

Tryptic digests were analyzed by LC-MS/MS on a QExactive HF mass spectrometer (Thermo Fisher Scientific) coupled with an Ultimate 3000. Peptides were separated by reverse phase (RP)-HPLC on a nanocapillary column, 75 μm id × 25cm 2μm PepMap Acclaim column. Mobile phase A consisted of 0.1% formic acid (Thermo) and mobile phase B of 0.1% formic acid/acetonitrile. Peptides were eluted into the mass spectrometer at 300 nL/min with each RP-LC run comprising a 90-minute gradient from 10-25% B in 65 min to 25-40% B in 25 min. The mass spectrometer was set to repetitively scan m/z from 300 to 1400 (R=240,000) followed by data-dependent MS/MS scans on the twenty most abundant ions, minimum AGC 1e4, dynamic exclusion with a repeat count of 1, repeat duration of 30s, (R=15000) FTMS full scan AGC target value was 3e6, while MSn AGC was 1e5, respectively. MSn injection time was 160 ms; microscans were set at one. Rejection of unassigned and 1+, 6-8 charge states was set.

### Data processing for proteomics analysis

Raw MS files were processed using MaxQuant, version 1.5.7.4 for identification of proteins (39). The peptide MS/MS spectra were searched against the UniProtKB/Swiss-Prot Mouse Reference Proteome database, (Proteome name, *Mus musculus* C57BL/6J – Reference proteome; Proteins, 49,838; Proteome ID, UP000000589; Strain, C57BL/6J; Taxonomy,10090 - *Mus musculus;* Last modified, July 9, 2016; Genome assembly, GCA_000001635.6). Precursor ion tolerance was 4.5 ppm with semi-tryptic specificity and MS2 fragment ion tolerance was set to 20 ppm. Oxidation of methionine, acetylation of the protein N-terminus and conversion of glutamine to pyroglutamic acid were used as variable modifications and carbamidomethylation of cysteine was set as a fixed modification. The minimal length required for a peptide was seven amino acids. Target-decoy approach was used to control false discovery rate (FDR). A maximum FDR of 1% at both the peptide and the protein level was allowed. The MaxQuant match-between-runs (0.7 min.) feature was enabled.

### Statistical Analysis

Statistical analysis was performed using GraphPad Prism. Data were expressed as mean+/− s.e.m. Two-tailed Student’s t-test was used for comparisons of two groups. A *P*-value of less than 0.05 was considered statistically significant.

## Acknowledgements

This study was supported by the National institute of Health (NIH: PO1 HL120846, PO1 HL40387 and K08 HL119597) and the American Society of Hematology Scholar Award (S.M.)

## Author Contributions

S.M. designed, conducted and analyzed experiments, co-supervised the project, and wrote the manuscript. A.S. conducted experiments and wrote the manuscript. L.W. conducted and analyzed some experiments and wrote the manuscript. J.G. conducted some experiments. L.Z. analyzed the data. F.G. conduced some experiments. L.A.S. conducted some experiments. S.H.S designed and analyzed some experiments. E.B. analyzed the data. C.S.A. analyzed the data, co-supervised the project and wrote the manuscript. All the authors reviewed the manuscript.

